# Tracing immune cells around biomaterials with spatial anchors during large-scale wound regeneration

**DOI:** 10.1101/2022.03.24.485669

**Authors:** Yang Yang, Chen-yu Chu, Yi Man, Li Liu, Chen-bing Wang, Yi-li Qu

## Abstract

Inevitable skin scarring devoid of dermal appendages bring about unfavorable effects on both aesthetic and physiological functions. Here we offered an available method for large-area wound regeneration using biodegradable Aligned extracellular matrix (ECM) scaffold. Accelerating wound coverage and enhanced hair follicle (HF) neogenesis were observed with the implantation of biomaterials. Multimodal analysis during wound regeneration provides a unique lens through which we can explore the foreign body responses (FBR) around biomaterials. First, we identified cell heterogeneity in responses to ECM scaffolds using single-cell RNA sequencing. Next the spatial anchor highlighted preferential recruitment of T cells around biomaterials colocalized with hair follicle precursor cells. Reconstruction of signaling pathways suggested the pro-regenerative effect of the adaptive immune systems on HF neogenesis. Immunodeficient mice lacking mature T lymphocytes lost the ability to activate papillary fibroblast as well as stimulate HF regeneration, validating the potential therapeutic effect of the adaptive immune system activated by biomaterials. In conclusion, this study provided the further understanding of the FBR and facilitate the design of immunoregulatory biomaterials for biomedical application in the future.

## Introduction

After severe skin damage, resulting scar usually contain dense extracellular matrix (ECM) fibers devoid of hair follicle (HF) and sebaceous gland (SG), which lack sensation and endocrine function as well as flexibility of normal skin^1^. As a result, there is an urgent need to explore the key mechanisms stimulating HF regeneration in skin repair.

In fact, tissue regeneration mediated by immunoregulatory biomaterials is emerging as a perspective strategy in tissue engineering^2,3^. Biological cues can be integrated into a polymer scaffold to mimic the native ECM, which guides tissue regeneration^4,5^. Upon materials implantation, the FBR process initiates with an immune response. It has been shown that modulating FBR by adjusting biomaterials characteristic may create a desired biological response to mobilize stem cells or stimulate specific cell proliferation ^4,5^. Previous study reported that the Aligned nanofibers scaffold with an immunomodulatory effect in accelerating small skin wound (diameter = 0.6cm) healing^6,7^. Since the delayed wound closure and susceptibility to infection in large scale defects^8^, the pro-regenerative effect of ECM scaffold in large wound needs further evaluation. What’s more, as immune cells are recruited into the scaffold to create a desirable microenvironment for cell adhesion, proliferation, and tissue reconstruction, the interactions between immune cells and cutaneous cell become more complex.

Currently, high resolution technique such as single-cell RNA sequencing (scRNA-seq)^6,9,10^ has been applied to identifies cell subpopulations in implantation model but lack the information of spatial distribution. Development in spatial transcriptomics (ST) has enabled the assessment of gene expression at spatial resolution^11^, which has been applied to the study of cancer^12 13^, liver^14^, and brain tissue^15^ to detect the regional cellular communication. The wound healing model with implanted biomaterials provide an ideal method to understand the FBR and probe the role of immune system in tissue regeneration. To our knowledge, multiomic approaches, including scRNA-seq and ST, were firstly applied to trace spatial heterogeneity in the biomaterial-mediated wound healing process in this study. Here we unveiled the cell composition around the ECM scaffold in both wildtype and immunodeficient mice to anchor the key role of T cells in HF regeneration, which would help optimize the existing biomaterial constructs and provide feasible strategies for the design of novel immunoregulatory products in tissue engineering for biomedical use.

## Results

### Enhanced HF neogenesis in scaffold-implanted wounds with activation of the adaptive immune system

To minimize wound contraction by the panniculus carnosus of rodents and induce scar large enough to observe^16^,we created a large area splinted full-thickness wound healing model in C57BL/6 mice to investigate whether a further regenerative phenomenon was observed in ECM scaffold-treated wounds. The workflow for evaluating large wound healing is summarized in Fig. 1a. We placed ECM scaffolds below the large wound (diameter = 1.5 cm) in the ECM_LW group, while the Ctrl_LW group received no biomaterials (Fig. 1b). Wound coverage was faster in the ECM_LW group on day 7 (Fig. 1 c, d), and the ECM_LW group also had a smaller gap width on day 7 and day 14 (Fig. 1e,f). Immunofluorescence (IF) staining for cytokeratin 5 (Krt5) and cytokeratin 10 (Krt10) showed that keratinocytes crawled around the biomaterials, and the ECM_LW group owned a larger area of neo-epithelium (Fig. 1h).

**Fig. 1.**
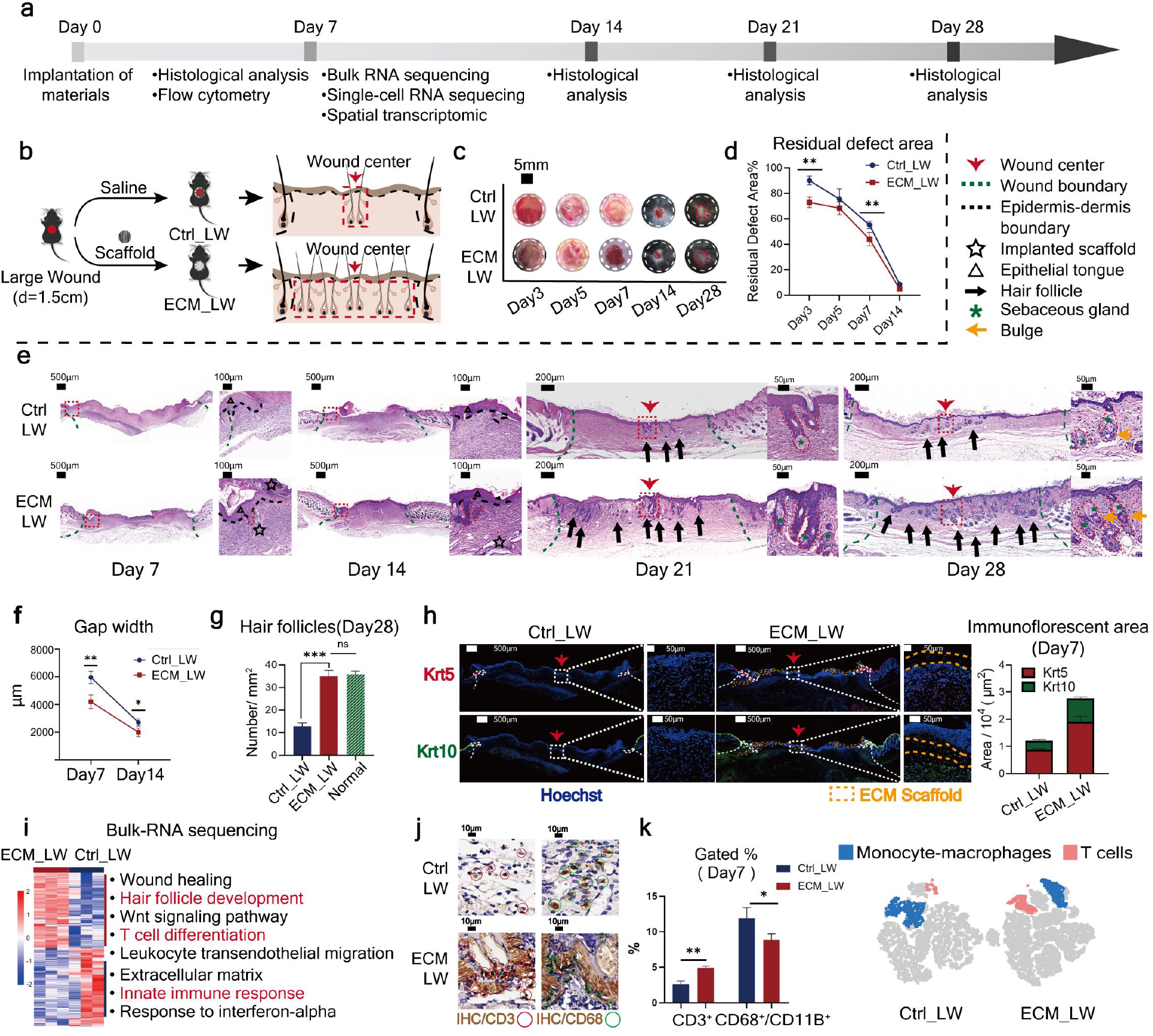
Evaluation of the wound healing process treated with ECM scaffolds. (a) Workflow for evaluating large-scale wound healing. (b) Surgical processes for skin excisional wound model. (c) Residual wound area at 3, 5, 7, 14 and 28 days, and (d) corresponding analysis (n=3 for each group). (e) H&E analysis of two groups at 7, 14, 21 and 28 days. (f) semiquantitative evaluation of gap width (n=3 for each group). (g) Histologic quantification of de novo HFs on POD28 (n=3 for each group).(h) IF staining using Krt5 (red) and Krt10 (green), and semiquantitative evaluation of the fluorescent area. (i) Heatmap showing hierarchical clustering of differentially expressed genes (p value<0.05 & |log2FC|> 1) and gene set enrichment analysis (n=3 for each group).(j) IHC staining of CD3 and CD68 at 7 days. (k) In vivo flow cytometry strategy using CD3 to identify T cells, CD68 and CD11b to identify monocyte-macrophages, and visualization of the cells in the tSNE (t-distributed Stochastic Neighbor Embedding) plot. ***P < 0.0001, **P < 0.01, and *P < 0.05 by Student’s t-test for data in (d), (f),(h) and (k), and analysis of variance (ANOVA) for data in (g).

Rare capability of adult mammals for limited HF regeneration^17,18^ was observed in Ctrl_LW group: there were limited HF neogenesis restricted in wound center since the POD21. In comparison, we observed that the ECM_LW group recapitulated the normal skin architecture with an equivalent number of mature HF on POD28 (Fig. 1g). The outline of HF morphogenesis could be recognized in Fig 1e, with the first appearance in the hair germ stage on POD7, they gradually grew into a bulbous-peg structure on POD21, and fully regenerated with mature hair shafts, dermal papillae, sebaceous glands, and bulges on POD28. Of note, the ECM membrane implanted in the large wound did not trigger any obvious fibrous capsule and exhibited an appropriate degradation rate in vivo. We observed degrading fragments on POD14, and there were no visible particles remaining on POD28 (Fig. 1e). The excellent biocompatibility and degradability of the membrane could prevent the risk of immune rejection and secondary surgery in further clinical applications.

To explore the underlying mechanism, Bulk-tissue RNA-seq (bulk-seq) analysis was conducted for two groups harvested on day 7 (n =3 for each group). We noticed that the ECM_LW group showed increased enrichment of genes involved in hair follicle development along with T cell differentiation (Fig. 1i). Immunohistochemistry (IHC) staining and flow cytometry confirmed increased T-cell infiltration from ECM_LW group in early phase (Fig. 1 j, k). In Ctrl_LW group, IF staining showed the accumulation of CD3-expressing T cells and Lgr5-espressing hair follicle stem cells (HFSC) limited in wound center on day7, day 14 and day 21 (Extended Data Fig. 1a-c), whereas biomaterials resulted in the robust recruitment of T cells and HFSC within and around the scaffold on day7 and day14 (Extended Data Fig. 1a-c).

### A single-cell atlas of the biomaterials tissue microenvironment

To explore the spatial characteristics of single cells during wound healing, we applied ST and scRNA-seq to compare the spatial gene expression profiles between two groups (Fig. 2a). At first, to explore the cell composition in biomaterial-treated wound, we isolated cells from the ECM_LW and Ctrl_LW samples and applied them to the 10x scRNA-seq platform (Extended Data Fig. 2a). After cell filtering, 31,740 single-cell transcriptomes were included in the final dataset (20,312 for the ECM_LW group and 11,428 for the Ctrl_LW group). Unsupervised clustering using Seurat categorized the cells into 10 main classes of cells. The composition of each cell cluster was listed so that the proportion of cells from two groups could be identified across all cell clusters (Extended Data Fig. 2b).

**Fig. 2.**
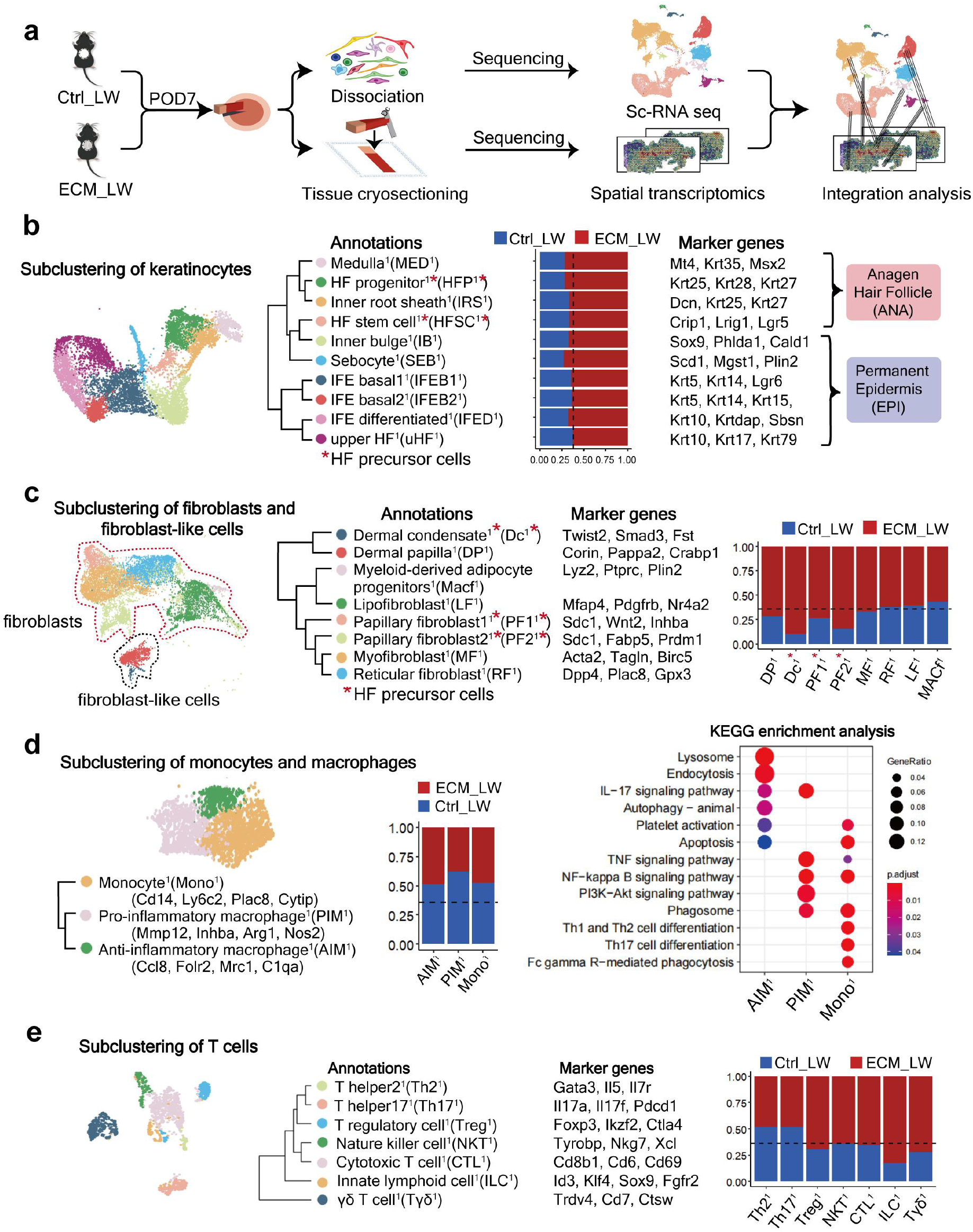
The single-cell atlas of biomaterials mediated microenvironment. (a) Schematic for generating scRNA-seq and spatial transcriptomics data from large area excisional wounds. (b) Subclustering of keratinocytes showing four subsets from anagen hair follicle and six subsets from permanent epidermis. The composition and marker genes for each subset are listed. (c) Subclustering of fibroblasts showing two fibroblast-like subsets and six fibroblast subsets. The marker genes and composition for each subset are listed. (d) Subclustering of monocyte/macrophage showing three subsets. The marker genes, composition and KEGG enrichment analysis for each subset are listed. (e) Subclustering of T cells showing seven subsets. The marker genes and composition for each subset are listed.

In previous report, HF neogenesis occurs through the migration of epithelial HFSC or hair follicle progenitor (HFP) to the wound center and form the placodes to activate papillary fibroblast (PF) fate specification into dermal condensate (Dc)^17,19,20^. The heterogeneity of keratinocyte and fibroblast subclusters of this dataset corresponded with the healing outcomes of two groups. Firstly, for subclustering analysis of keratinocyte (Fig. 2b), the higher proportion of Krt5^+^Krt14^+^ interfollicular epidermal basal cell^1^ (IFEB^1^) and Krt10^+^Krtdap^+^ interfollicular epidermal differentiated cell^1^ (IFED^1^) were observed in the ECM_LW group, supporting more neo-epithelium proliferation in the presence of scaffolds. In addition, we found the higher proportion of Krt25^+^Krt28^+^Krt27^+^ hair follicle progenitor^1^ (HFP^1^) ^6^ and Crip1^+^Lrig1^+^Lgr5^+^ hair follicle stem cell^1^ (HFSC^1^)^21^ in ECM_LW group, which served enough epithelial resources for HF reconstruction. Next, we defined two fibroblasts and six fibroblast-like cells subclusters based on defined markers published before^6,19,22,23^ (Fig. 2c). In accordance with previous report^19,24^, the major initial wave of dermal repair in Ctrl_LW group was mediated by the lower lineage fibroblast including Mfap4^+^ Cd34^+^ lipo-fibroblast^1^ (LF^1^) and Dpp4^+^Mgp^+^Gpx3^+^Plac8^+^ reticular fibroblast^1^ (RF^1^), which was respectively related to hypodermis adipocytes formation and dermis collagen fibril organization in gene ontology (GO) enrichment analysis (Extended Data Fig. 3a). In contrast, there were more upper lineage PF in biomaterial-implanted wound. The PF in this dataset consisted of 2 subclusters including PF1^1^ (Runx1+Inhba+)^22^ and PF2^1^ (Notch1+Prdm1+)^25^, both of which were believed to have the capacity to support HF initiation (Extended Data Fig. 3a). For fibroblast-like subclusters, we observed higher proportion of (Papp2^+^Corin^+^) dermal papilla^1^ (DP^1^) cells in ECM_LW group, which resided in the mature HF base in normal skin. Remarkably, we identified the Smad3^+^Lef1^+^Krt17^+^ dermal condensate^1^ (Dc^1^) cell^20^ in this dataset. Derived from PF, the Dc was believed to be the origin of DP^20,26^. In embryologic HF morphology, Dc is acting as the signaling niches to stimulate epithelial placode growth^20^, and thus promote HF morphogenesis. Since approximate 90% of Dc^1^ were from ECM_LW group, as well as significantly higher proportion of PF1^1^ and PF2^1^, which might provide enough mesenchymal component for the subsequent HF formation in biomaterial implanted group too.

For Ptprc+ immune micro-environment, most myeloid cells owned the lower proportion in ECM_LW group (Extended Data Fig. 2b). Firstly, as the first responding immune cells acting as the critical mediators of innate immune system, neutrophils are necessary for the recruitment and differentiation of monocytes^27^. Four neutrophil subsets including Il1b^+^Retnlg^+^ peripheral blood neutrophil^1^ (PBN^1^), Il1a^+^Tnf^+^ neutrophil1^1^ (Neu1^1^), Lrg1^+^Il1r2^+^ neutrophil2^1^ (Neu2^1^), Ccl9^+^Rgs1^+^ neutrophil3^1^ (Neu3^1^) were identified in our dataset (Extended Data Fig. 4a). Neu3^1^ was the only subsets owned a higher proportion in ECM_LW group and showed gene enrichment in leukocyte migration and Th17 cell differentiation. Dendritic cell (DC) was classified into two subclusters including Krt5^+^Krt14^+^ langerhans cell^1^ (LC^1^) and Cd86^+^ monocyte derived dendritic cell^1^ (MDC^1^) (Extended Data Fig. 4b). Kyoto Encyclopedia of Genes and Genomes (KEGG) enrichment analysis showed that LC^1^ and MDC^1^ respectively related to phagosome and antigen presentation. There was no obvious difference in the proportion of DC subsets between the two groups. Monocytes and macrophages were of vital importance in orchestrating biomaterial related FBR and tissue regeneration^28^. Three subsets of macrophages including Ccl8^+^Folr2^+^C1qa^+^Mrc1^+^ anti-inflammatory macrophages^1^ (AIM^1^), Mmp12^+^Arg1^+^Inhba^+^Nos2^+^ pro-inflammatory macrophages^1^ (PIM^1^) and Ly6c2^+^Plac8^+^Cytip^+^H2-Eb1^+^ monocytes^1^ (Mono^1^) were determined. The ECM_LW group contributed to a lower proportion of all three subsets, suggesting that the scaffold might play a role in the reduction of macrophage infiltration at the proliferative stage (Fig. 2d).

T cells were the only immune cell which owned a higher proportion in ECM_LW group (Extended Data Fig. 2b), and genes related to T cells (Il7r, Areg and Tubb5) were substantially higher expressed by the ECM_LW group (Extended Data Fig. 2d), suggesting the activation of adaptive immune system induced by biomaterials. Subclustering of T cells resulted in seven subsets including Cd8b1^+^ cytotoxic T cell^1^ (CTL^1^), Tyrobp^+^Nkg7^+^Xcl^+^ nature killer cell^1^ (NKT^1^), Il17a^+^Il17f^+^ T helper17^1^ (Th17^1^), Foxp3^+^ T regulatory cell^1^ (Treg^1^), Trdv4^+^Cd7^+^Ctsw^+^ γδT cell^1^ (Tγδ^1^), Gata3^+^Il5^+^ T helper2^1^ (Th2^1^) and Bcl11a^+^Id3^+^Notch1^+^ innate lymphoid cell^1^ (ILC^1^) based on markers from published research^29,30^. Derived from common lymphoid progenitors, ILCs mirror αβ T cell subtypes functionality responding to local signals but lack antigen-specific receptors. In skin, tissue resident ILCs play key roles in barrier integrity, inflammatory response initiation, and immunological response modulation in wound repairing^30-33^. Correlation analysis showed that ILC^1^ was highly correlated with CTL^1^ (Extended Data Fig. 3c). Remarkably, skin-resident ILC^1^ in our dataset possessed a tissue specific transcriptional profile that co-expression genes related to T cell functionality (Cd3d^+^Klf4^+^Fst^+^Itga6^+^) and HF morphology (Wnt4^+^Sox9^+^Fgfr2^+^). Gene enrichment analysis showed the ILC was enriched in bacterial invasion of epithelial, leukocyte transendothelial migration as well as Wnt signaling pathway (required in hair follicle regeneration^17^) (Extended Data Fig. 3b), suggesting that ILC^1^ might initiate HF neogenesis by balancing reparative and pro-inflammatory functions^34^.

### Discovery of spatial heterogeneity around biomaterials

To explore the spatial characteristics of cell heterogeneity in scaffold-implanted wounds, we applied ST analyze the spatial gene expression profiles. Anatomical structures were identified in the ST sections (Fig. 3a). Based on the gene enrichment analysis of ST, the ECM_LW sample showed up-regulated gene expression in hair follicle morphogenesis as well as adaptive immune response (Fig. 3b). The spatial feature plot illustrated keratinocytes crawling around the implanted biomaterials (Fig. 3c). Moreover, the distribution of PF^1^ (HF-related) was enlarged in the ECM_LW group (Fig. 3d). In contrast, lower lineage fibroblasts RF^1^, were significantly reduced, suggesting different healing outcomes of two groups. Generally, HF formation does not start until several days after wound closure (POD14–21) in large wounds without implantation^17^. In the ST profiling of the ECM_LW group on POD7, we identified the initial structure of the HF composed of Krt28^+^ HFP in the neo-epidermis and Twis2^+^ Dc aggregation beneath the dermis (Fig. 3e), which highlighted the early HF formation signature in the ECM_LW group at the spatial gene expression level (Fig. 3f).

**Fig. 3.**
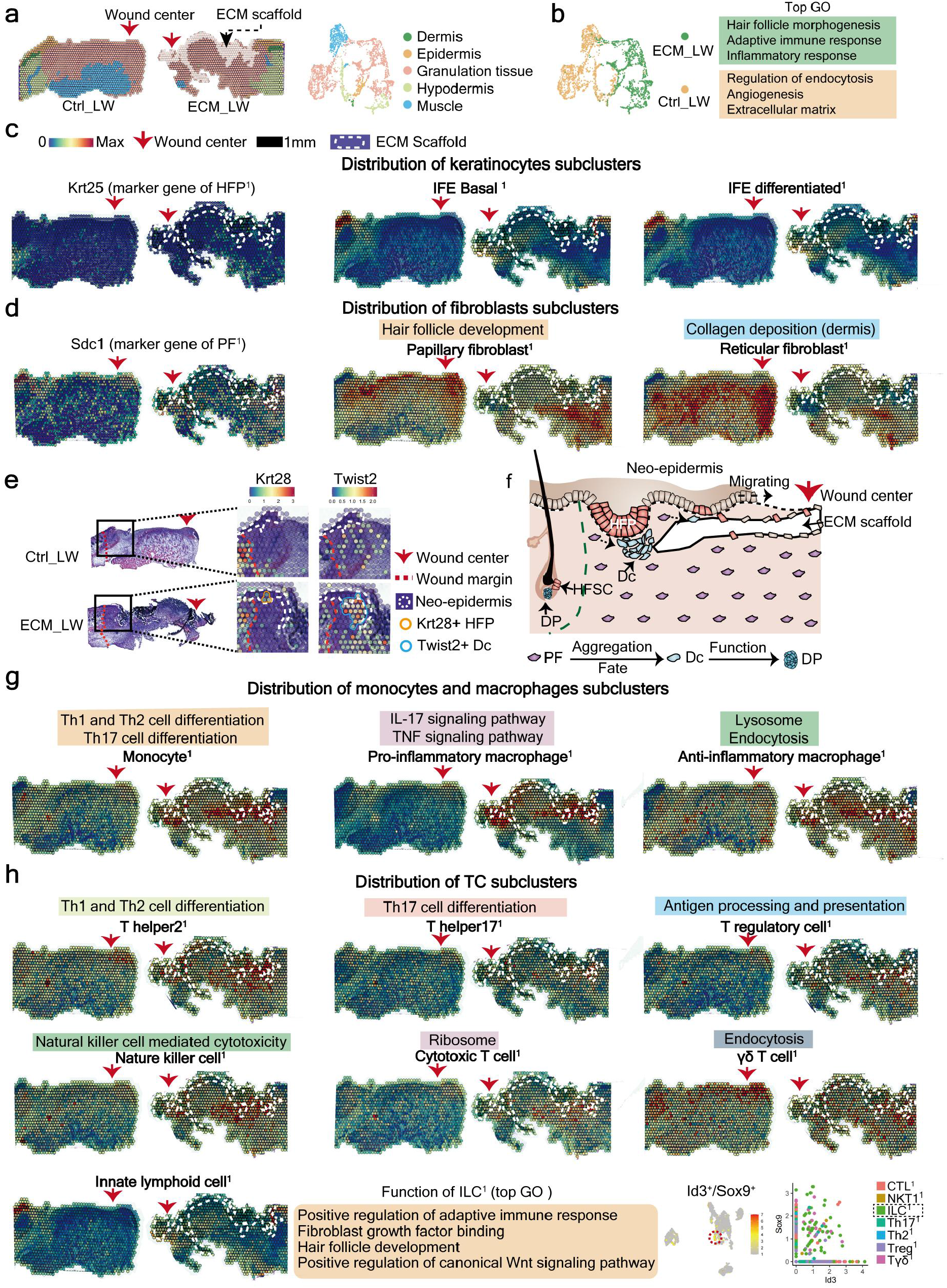
Spatial anchors tracing the cell distribution around the ECM scaffold. (a) The unsupervised clustering indicated the anatomical structure of samples. (b) Gene enrichment analysis between ECM_LW and Ctrl_LW group. (c) Spatial feature plot showing the expression of Krt25 (marker gene for hair follicle progenitor) and defined keratinocyte subclusters in tissue sections. (d) Spatial feature plot showing the expression of Sdc1 (marker gene for papillary fibroblast) and defined fibroblast subclusters in tissue sections. (d) Spatial feature plot highlighted the expression of Krt28^+^ hair follicle progenitor and Twist2^+^ dermal condensate in migrating neo-epidermis. (f) Illustration showing the epithelialization along with de novo HF formation in biomaterials mediated healing process. (g) Spatial feature plot showing the expression of defined monocyte, macrophage (MAC) subclusters in tissue sections. (h) Spatial feature plot showing the expression of defined T cell subclusters in tissue sections.

To explore the scaffold-induced immune microenvironment, we integrated marker genes of immune cells (defined in Fig. 2) with ST profiling. We noticed the obvious aggregation of monocyte-macrophage (MAC) and T cell subclusters surrounding the biomaterial (Fig. 3g,h). Notably, we observed the accumulation of tissue-resident ILC^1^ in the epithelium and granulation tissue in the ECM_LW group (Fig. 3h). ILCs that co-expressed genes related to the adaptive immune response as well as hair follicle development may be critical regulators initiating HF formation.

### Prediction of critical signaling patterns between spatially co-localized cell populations around the biomaterial

Based on the multimodal profiling, we observed the higher proportion (Fig. 4a) and overlapping distribution of T cells and HF precursor cells (Fig. 4b) in the ECM_LW group, which implied a potential cellular communication among them. Next, we applied CellChat to predict the cell-cell communication patterns between EMC_LW and Ctrl_LW. The heatmap shows the overall signaling strength of the ligand-receptors in the Ctrl_LW and ECM_LW groups (Fig. 4c). Adaptive immune system-related pathways, including MHC-II, were further activated in ECM_LW. In addition to IL6, key signaling pathways in embryonic HF development such as fibroblast growth factor (FGF) ^20^, NOTCH ^35^, and KIT ^36^ were also upregulated in the ECM_LW group. The critical functions of NOTCH ^35^, pleiotrophin (PTN), and midkine (MK) ^36^ in skin morphology have been emphasized previously. Consistent with the gene expression profile in single cell level, adaptive immune cells and HF precursor cells were enriched with interacting receptor/ligand gene partners, including Jag1-Notch2, Mdk-(Sdc1 or Itgb1), and Ptn-(Itgb1or Sdc1) in ST profiles (Fig. 4 d, e). Based on analysis above, adaptive immune cells might modulate HF precursor cell growth through these pathways, thus facilitating HF formation.

**Fig. 4.**
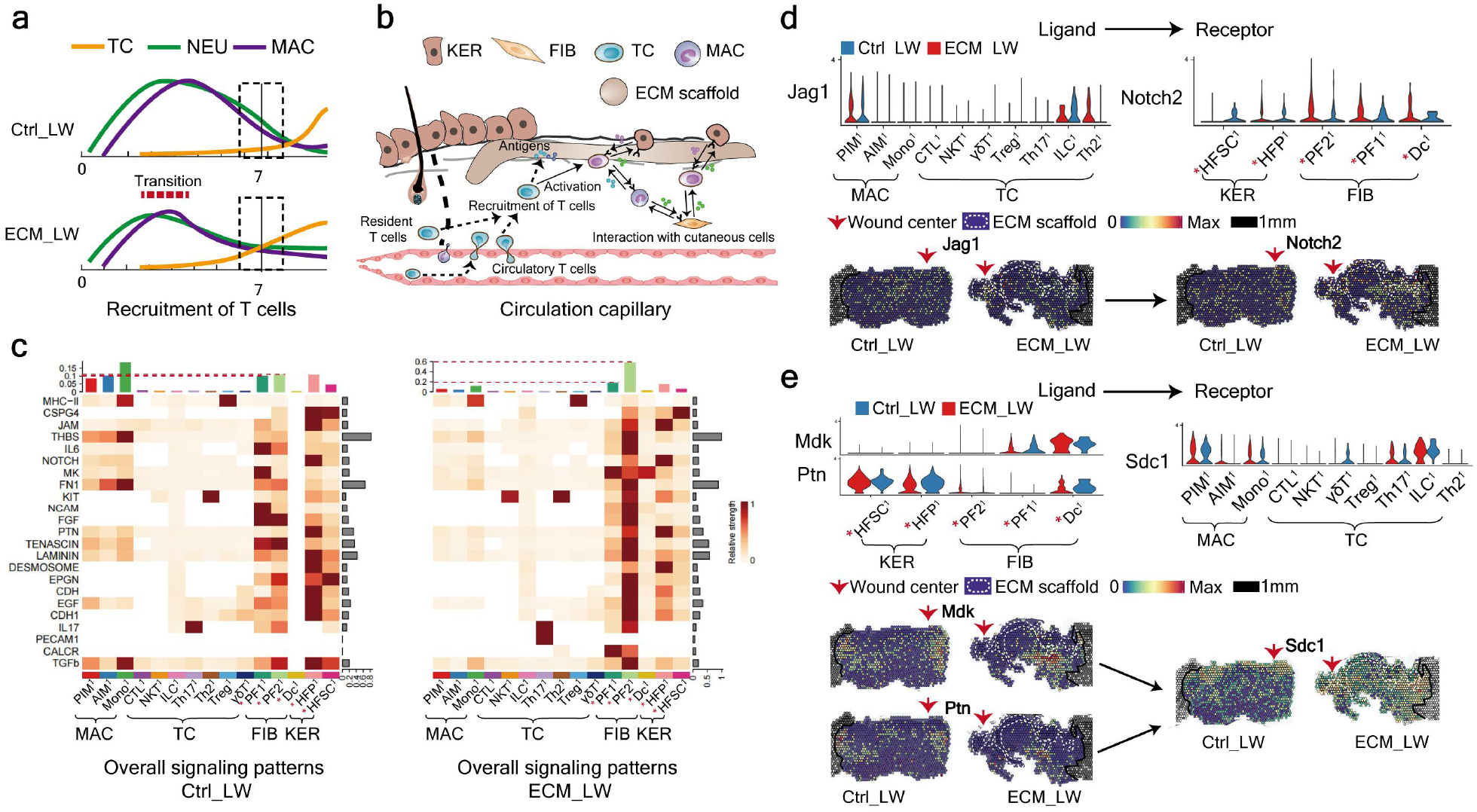
Cellular communication landscape between immune cells and cutaneous cells. (a) Schematic timeline highlighting the recruitment of immune cell from innate and adaptive immune systems in Ctrl_LW (top) and ECM_LW (bottom) groups. (b) Schematic of combined ligand-receptor analysis between cutaneous cells and adaptive immune cells in biomaterial treated group. (c) Comparison of overall cell-cell communication patterns between Ctrl_LW and ECM_LW using CellChat. (d) Violin plot (scRNA-seq) and spatial feature plots (ST) showing select ligands and cognate receptor expression of NOTCH signaling pathway. (e) Violin plot (scRNA-seq) and spatial feature plots (ST) showing select ligands and cognate receptor expression of MK and PTN signaling pathways.

### Adaptive immune system was required for the skin regeneration mediated by ECM scaffold

To determine the role of adaptive immune system in HF regeneration, we placed ECM scaffolds in the dorsal skin of immunodeficient C57Bl/6 (B6.129-Rag2tm1) mice, which lacked mature T lymphocytes and innate lymphoid cells ^37,38^ (Fig. 5 a, b). We noticed that the wound closure rate was delayed on day7 (Fig. 5c-5e), and regeneration of HF was scarce (Fig. 5c, f) on day 28 in Rag2^−/−^ mice. Compared to wild type (WT) mice, Rag2^−/−^ group still possess a myeloid recruitment ability, but no visible accumulation of T cell around biomaterials was observed (Fig. 5 g, h). Next, we further explored the differences in cell composition and spatial gene expression between WT and Rag2^-/-^ mice using scRNA-seq (Extended Data Fig. 5a). Unsupervised clustering categorized the cells into nine clusters based on gene expression (Extended Data Fig. 5 b, c). Firstly, subclustering analysis of mesenchymal cell identified two fibroblast-like cell and five fibroblast cell subsets (Fig. 5i). In accordance with the HF regeneration outcomes, the Rag2^-/-^ group reduce the proportion of PF^2^ (related to HF development) but improve the number of RF^2^ (related to collagen deposition) and LF^2^ (related to collagen deposition and angiogenesis). For immune microenvironment, the Rag2^-/-^ samples recruited less adaptive immune cells (Cd3g^+^Cd7^+^ T cells) and more innate immune cells (Lyz2^+^C1qb^+^ monocytes and macrophages) than the WT group (Extended Data Fig. 5b). T cell related genes such as Areg, Trdc and Rgs2 were down-regulated in Rag2^-/-^ samples (Extended Data Fig. 5d). The proportion of neutrophil and dendritic cell subsets was basically equilibrium between the two groups (Extended Data Fig. 6a,b). Subclustering of macrophages resulted in four subsets, which were PIM^2^, AIM^2^, Mono^2^, and MDC^2^ (Fig. 5j). Rag2^-/-^ samples showed the lower number of Mono^2^ (related to T helper cell differentiation), but improve the recruitment of PIM^2^, AIM^2^, and MDC^2^. Subclustering of T cells resulted in seven subsets (Fig. 5k), the Rag2^-/-^ groups contributed to less T cells in most subsets except NKT^2^ (related to natural killer cell mediated cytotoxicity) and Th2^2^, indicating the reduction of adaptive immune systems in Rag2^-/-^ group.

**Fig. 5.**
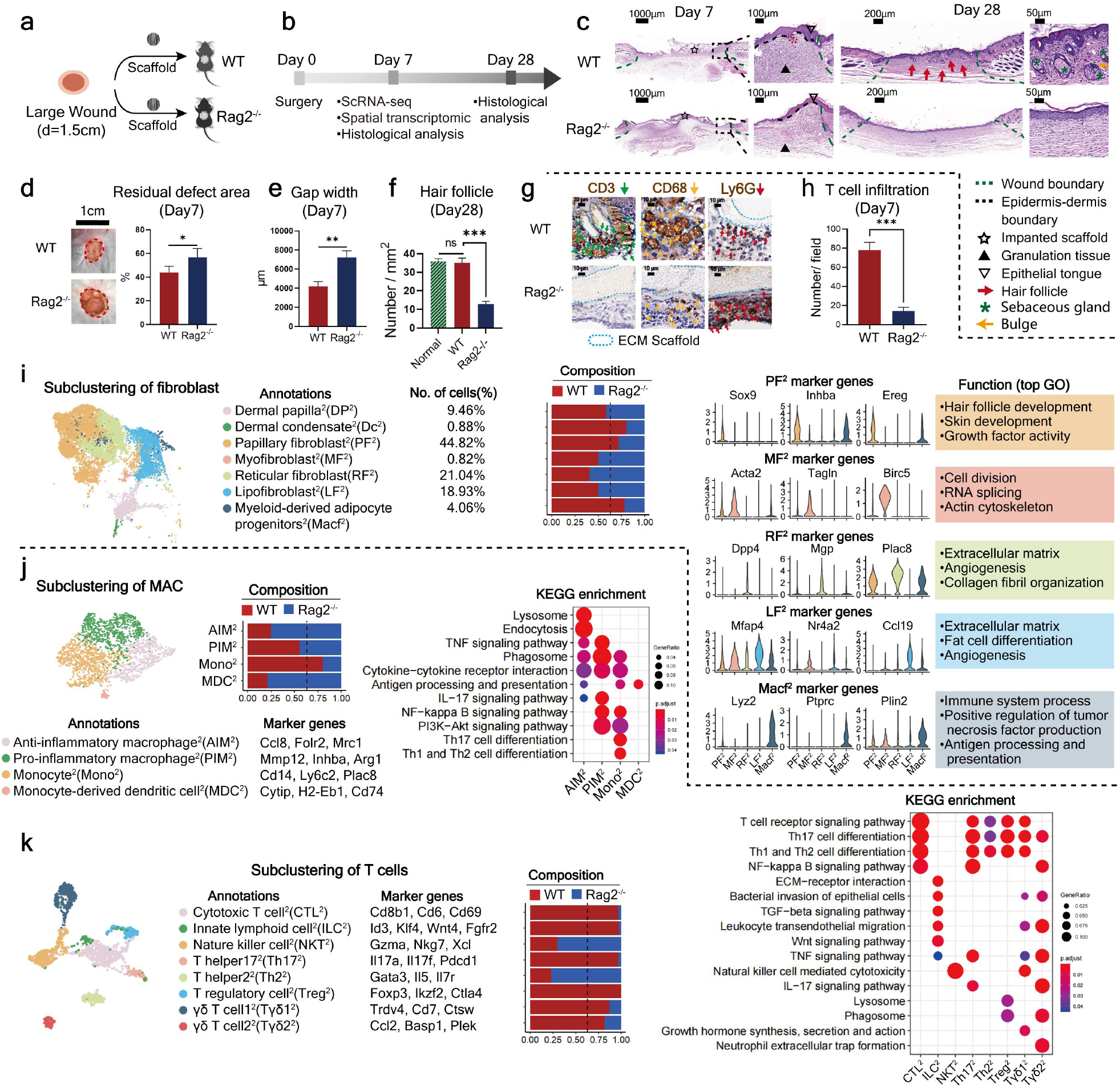
Evaluation of wound healing in immunodeficient mice lacking mature T cells. (a) Surgical process and (b) workflow for evaluating large area wound healing mediated by ECM scaffold in WT and Rag2^−/−^ mice. (c) Histological analysis of wound healing in WT and Rag2^−/−^ mice at 7 and 28 days. (d) Residual defect area at 7 days after wound (n=3 for each group). (e) Semiquantitative evaluation of gap width (n=3 for each group). (f) Histologic quantification of de novo HFs (n=3 for each group). (g) IHC staining of CD3 and CD68 and (h) semi-quantification of CD3^+^ T cells infiltration (n=3 for each group). (i) Subclustering of fibroblasts showing two fibroblast-like subsets and five fibroblast subsets. Marker genes for fibroblast subsets and their enriched gene sets in GO analysis are listed. (j) Subclustering of monocyte/macrophage showing four subsets. The marker genes, composition and KEGG enrichment analysis for each subset are listed. (e) Subclustering of T cells showing eight subsets. The marker genes, composition and KEGG enrichment analysis for each subset are listed. ***P < 0.0001, **P < 0.01, and *P < 0.05 by Student’s t-test for data in (d), (e) and (h), and analysis of variance (ANOVA) for data in (f).

To explore the spatial characteristics of cell heterogeneity in immunodeficient mice, we also applied ST analyze the spatial gene expression profiles. Anatomical structures could be identified in the ST sections (Fig. 6a). Based on the gene enrichment analysis of ST, the Rag2^-/-^ sample down-regulated gene expression in hair follicle morphogenesis but improve the genes related to collagen fibril organization (Fig. 6b). Compared to WT samples, the distribution of PF^2^ (HF-related) was reduced. Lower lineage fibroblasts RF^2^ and LF^2^ were significantly enlarged instead, corresponding with the healing outcome of Rag2^-/-^ groups. To compare the scaffold-induced immune microenvironment, we integrated marker genes of immune cells (defined in Fig. 5 j, k) with ST profiling. We noticed the aggregation of monocyte-macrophages (MAC) surrounding the biomaterial in both group (Fig. 6d). The expression of the T cell subsets were dramatic declined(Fig. 6e), indicating the reduction of adaptive immune system in Rag2^-/-^ group

**Fig. 6.**
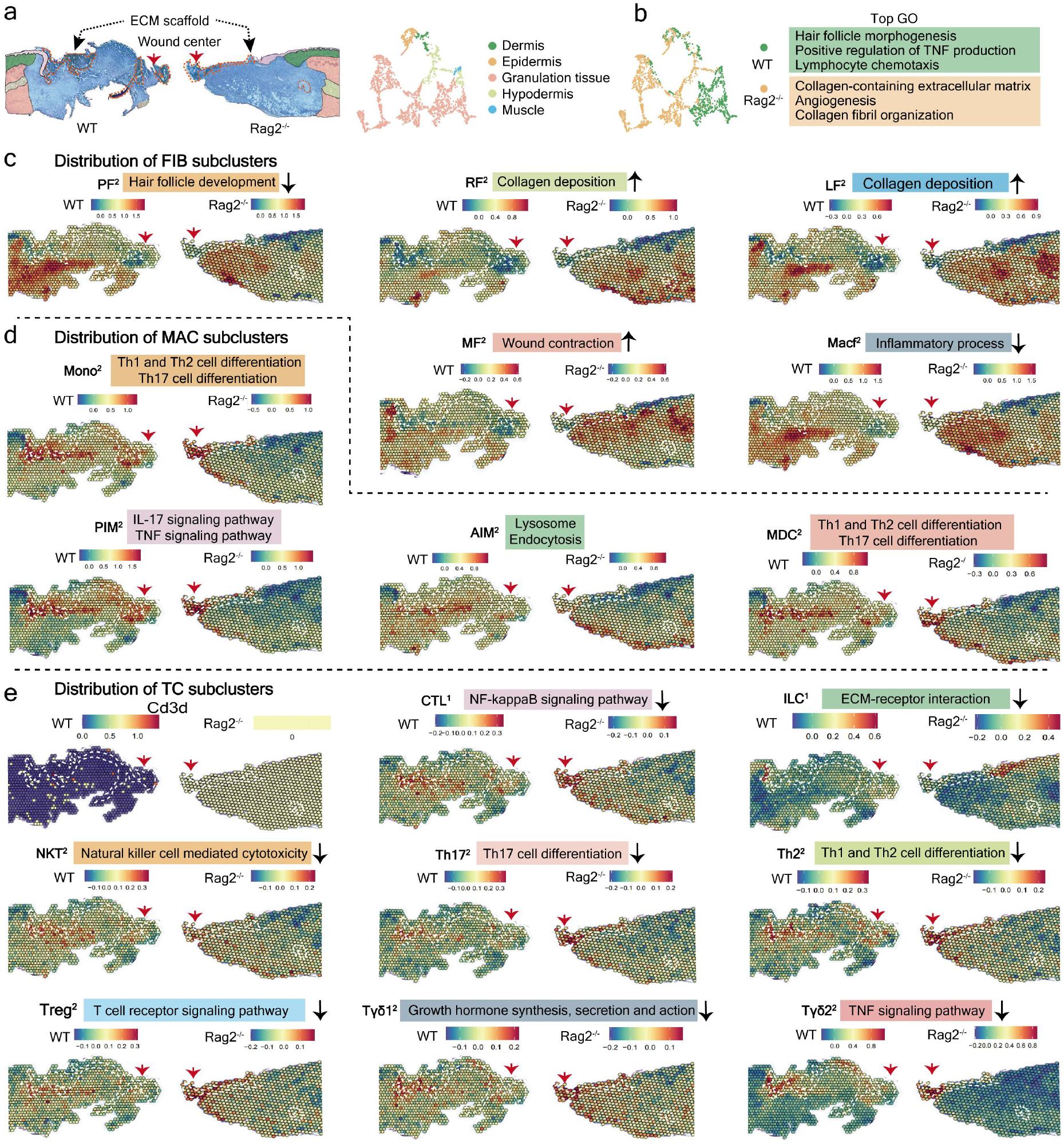
Spatial atlas of cell microenvironment around biomaterials of immunodeficient mice. (a) The anatomical structure of each sample which could be distinguished by Seurat. (b) Gene enrichment analysis between WT and Rag2^-/-^ group. (c) Spatial feature plot showing the defined fibroblast subclusters in tissue sections. (d) Spatial feature plot showing the expression of defined monocyte, macrophage (MAC) subclusters in tissue sections. (e) Spatial feature plot showing the expression of defined T cell subclusters in tissue sections.

### Biomaterials facilitate de novo HF regeneration despite of wound size

To detect any differences in wound healing between small and large wounds treated with biomaterials, we repeated the experiment with small (diameter=0.6 cm) and large (diameter=1.5 cm) full-thickness wounds implanted with biomaterials (Fig. 7 a, b). Consistent with the ECM_LW group, the scaffold implanted in the small wound (ECM_SW) did not trigger an obvious FBR fibrous capsule and had a rapid degradation rate (Fig. 7 c, d). Histological sections of ECM_SW tissue revealed clear signs of enhanced HF reconstruction too (Fig. 7 c, f). Based on bulk-seq profiling (n=3 for each group), heatmaps showed insignificant changes in HF-related, adaptive immune response, or innate immune response gene expression between the two groups (Extended Data Fig. 7a). IHC staining for CD3 and CD68 confirmed similar infiltration levels of T cells and monocyte-macrophages surrounding the scaffold (Fig. 7e).

We further explored the differences in immune cell composition using scRNA-seq and ST (Fig. 7 g, h). Feature plot showed the aggregation of Ptprc+ immune cells (Fig. 7i). The total proportion of MAC was in equilibrium between the two groups (Extended Data Fig. 7b), and integrated analysis of ST profiling showed similar distributions of the three MAC subsets around the scaffold (Fig. 7 j, k). Integrated analysis confirmed the substantial distribution of T cell subsets surrounding the scaffold in both the groups (Fig.7 l, m). In summary, the biomaterial showed similar “pro-regenerative” effect on wound healing despite the wound size. In addition, this resemblance to the immunoregulatory effect of the adaptive immune system indicates a significant role of modulating the immune environment in HF regeneration.

**Fig. 7.**
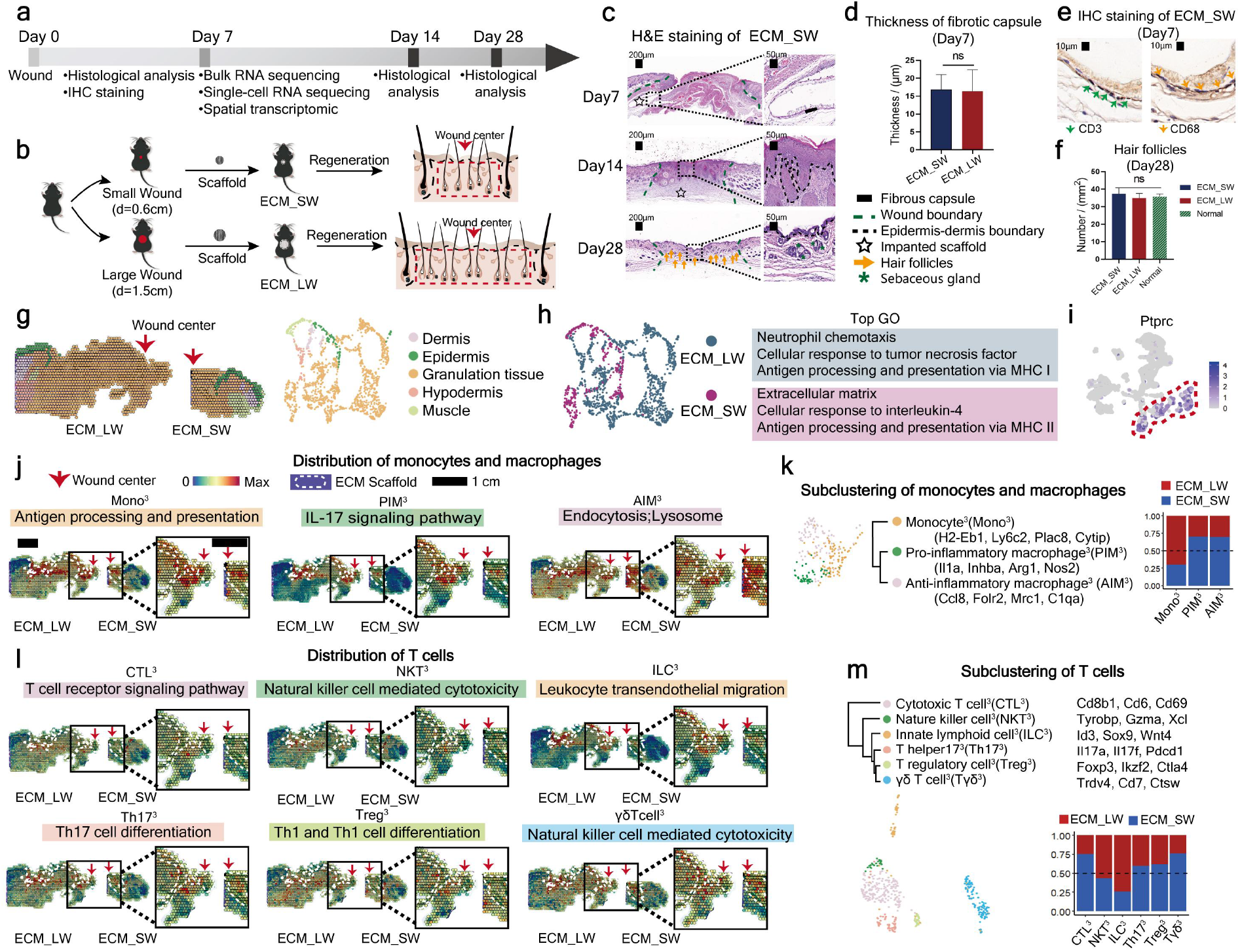
Evaluation of the healing of small and large full-thickness wound treated with ECM scaffolds. (a) Workflow for evaluating skin wound healing. (b) Surgical process for skin excisional wound model of ECM_SW and ECM_LW group. (c) H&E staining of ECM_SW samples. (d) Thickness of fibrotic capsule around membrane of ECM_SW and ECM_LW group (n=3 for each group). (e) IHC staining of CD3 and CD68. (f) Histologic quantification of de novo hair follicles on POD28 (n=3 for each group). (g) The unsupervised clustering indicated the anatomical structure of sample. (h) Gene enrichment analysis between ECM_LW and ECM_SW. (i) Feature plot showing the expression of Ptprc (Cd45) in scRNA-seq dataset. (j) Spatial feature plot showing the expression of defined monocyte, macrophage (MAC) subclusters in tissue sections. (k) Subclustering of MAC in sc-RNA seq dataset. (l) Spatial feature plot showing the expression of defined T cell subclusters in tissue sections. (m) Subclustering of T cell in sc-RNA seq dataset. ***P < 0.0001, **P < 0.01, and *P < 0.05 by Student’s t-test for data in (d), and analysis of variance (ANOVA) for data in (f).

## Discussion

Synthetic graft transplantation is an efficient treatment for severe large-area skin wounds, especially when the donor site is not qualified ^2^. With additional understanding of tissue engineering, it is suggested that synthetic biomaterials imitating the native structure of ECM can integrate and potentially play a pro-regenerative role in wound healing ^4,39^.The reconstruction of dermal appendages is an important indicator of full skin regeneration. Nevertheless, existing treatment can only form epidermal or dermal layers and fail to regenerate enough HF and SG^8^. Requirements for promotion of nascent HF formation bring significant challenges to biomaterial design, particularly for large-scale severe wounds.

Here, the pro-regenerative influence of the adaptive immune system in coordinating the early stages of wound repair has been stressed ^40^. We employed an ECM scaffold for large-area skin defects and investigated its immunoregulatory mechanism in wound healing. The scaffold showed an impact by accelerating wound closure and promoting nascent HF formation (Fig. 1 e, g). By multimodal analysis, we observed that substantial T cell infiltration in response to scaffold implantation was driven towards regenerative subpopulations (Fig. 2e, 3h), overlapping with the abundant PF (related to HF regeneration) (Fig. 3d). The local intercellular communications among the enlarged “pro-regenerative” immune niche and HF precursor cells suggested the essential role of the adaptive system in biomaterial-mediated wound regeneration (Fig. 4b). We next confirmed the requirement of T cells in skin regeneration by an immunodeficient model (Fig. 5). These data validated that HF can be reconstructed through the activation adaptive immune by ECM scaffolds. Meanwhile, mild FBR around the scaffold decreased the risk of immune rejection, and the appropriate degradation rate could also avoid the extra expense of secondary surgery in both small and large area wound healing (Fig. 7).

In this study, we offered an available manner for large-area wound regeneration and firstly defined the spatial heterogeneity of microenvironment during biomaterial-mediated wound healing process. These techniques provide a unique medium through which we can further understand the immunoregulatory mechanisms of the ECM scaffold. Of note, we testified the role of adaptive immune system activated by biomaterials in HF reconstruction and provides further insights into the future design of targeted immunoregulatory materials for scarless wound regeneration.

## Online Methods

### Fabrication of ECM scaffolds

The preparation of ECM scaffolds was similar to previous report^7^.To fabricate the electrospinning scaffold with approximate 300nm diameter aligned fibers, 20% w/v PLGA (LA/GA = 75:25, molecular weight = 105 kDa, Jinan Daigang Biomaterial Co. Ltd.) and 2% w/v FC (Sangon Biotech Co. Ltd.) were dissolved in HFIP (Aladdin Co., Ltd.) with stirring until complete dissolution. The solution was fed at 0.018mm/min, and a voltage of −2/5kv and 11 cm distance between the needle (21G) and the rotating cylinder (2800 rpm) was applied. After been dried in a vacuum oven for a week, The morphology of PLGA/FC nanofibrous scaffold was observed by the scanning electron microscopy (SEM; JSM-7500F, JEOL, Japan). The scaffolds were cut into circular shape (0.8cm or 1.8cm in diameter) and sterilized using γ-irradiation before the animal implantation experiments.

### Excisional wound model and implantation procedures

All procedures involving animals were approved by the Institution Review Board of West China Hospital of Stomatology (No. WCHSIRB-D-2017-033-R1). C57BL/6J mice (Dossy Experimental Animals Co., Ltd.) and immunodeficient C57Bl/6 (B6.129-Rag2tm1) mice (Shanghai Model Organisms Center Inc., Shanghai, China), at the age of 6 week (∼20 g) were used in this research. To minimize wound contraction by the panniculus carnosus of rodents and allow wound healing through the granulation and re-epithelialization like the human skin, we use the mice splinted model^16^. The circular (diameter =0.6 or 1.5cm) full-thickness wound were created in the mice dorsal skin and stented by silicone loops. The mice in further study were divided into three groups: large wound (diameter = 1.5cm) treated with saline (Ctrl_LW) or ECM scaffolds below the wound (ECM_LW), and small wound (diameter = 0.6cm) treated with ECM scaffolds (ECM_SW). Subsequently, the wounds were covered with sterile Tegaderm film (3M) and respectively fixed on the small (inner diameter 8 mm and outer diameter 12 mm, for small wound) or large (inner diameter 18 mm and outer diameter 22 mm, for large wound) silicone ring with uniform suture. Mice were euthanatized at 1, 2, 3, and 4 weeks after the surgery, and the small (diameter 10 mm, for small wound) or large (diameter 25 mm, for large wound) round full-thickness sample was harvested.

### Bulk RNA sequencing

Three replicates of mice skin wounds in each group were collected for the of bulk-tissue RNA sequencing procedure. Total amounts and integrity of RNA were assessed using the RNA Nano 6000 Assay Kit of the Bioanalyzer 2100 system (Agilent Technologies, CA, USA). mRNA was purified from total RNA by using poly-T oligo-attached magnetic beads. The library fragments were purified with AMPure XP system (Beckman Coulter, Beverly, USA). The PCR amplification product was purified by AMPure XP beads. After the library is qualified by qRT-PCR, the different libraries are pooling and being sequenced by the Illumina NovaSeq 6000. FeatureCounts (v1.5.0-p3) was used to count the reads numbers mapped to each gene. And then FPKM of each gene was calculated based on the length of the gene and reads count mapped to this gene. Differential expression analysis of two groups (three biological replicates per group) was performed using the DESeq2 R package (1.20.0), p−value <0.05 and |log_2_(foldchange)| >1 was set as the threshold for significantly differential expression. GO terms and KEGG pathways with corrected P-value less than 0.05 were considered significantly enriched by differential expressed genes.

### Single cell RNA sequencing

Specimen harvest procedure for scRNA-seq was like previous report^6^,3∼4 fresh samples were collected per group. In brief, the wound tissues were firstly digested by the Epidermis Dissociation Kit (Epidermis Dissociation Kit, mouse; Miltenyi Biotec) for enzymatic epidermal-dermal separation. The epidermis part was dissociated by a gentleMACS Dissociator (Miltenyi), then filtered (70-mm cell strainer, Corning, Durham), centrifuged (300g, 10 min, 4°C), and resuspended with phosphate-buffered saline (PBS) containing 0.5% bovine serum albumin (BSA). The dermis part was cut into 1 mm width pieces and mixed with mixed enzyme solution containing type I collagenase (Gibco, Grand Island) and trypsin (Gibco, Canada), then dissociated by gentleMACS Dissociator (Miltenyi), and digested for 2.5 hours in a hybridization oven (Peqlab PerfectBlot). After being dissociated, filtered, centrifuged, and resuspended in red blood cell lysis buffer (Solarbio), the dermis cells were mixed with epidermis part. Then the dead cells and debris were removed by Dead Cell Removal MicroBeads (Miltenyi). Lastly, these suspensions were then carried out for Single-Cell RNA-seq (10x Genomics Chromium Single Cell Kit). Sequencing (10x Genomics Chromium Single Cell 3′ v3) was performed using an Illumina 1.9 mode. Then, reads were aligned, and expression matrices were generated for downstream analysis (Cell Ranger pipeline software). Filtering, normalization, scaling and canonical correlation analysis were performed (Seurat 4.1.2.) for integrated datasets. CellChat was used to predict receptor-ligand probability among cell subpopulations.

### Spatial Transcriptomics

Fresh samples were rapidly harvested and frozen in OCT. Wound tissues were cryosectioned at - 20 degrees onto gene expression slides. Tissue sections on the Capture Areas of the Visium Spatial Gene Expression are fixed using methanol, H&E staining images will be used downstream to map the gene expression patterns. Using the Visium Tissue Optimization Slide & Reagent kit, permeabilization time was optimized. Second Strand Mix is added to the tissue sections to initiate second strand synthesis. After transfer of cDNA from the slide, spatially barcoded, full-length cDNA is amplified via PCR to generate sufficient mass for library construction. The final libraries contain the P5 and P7 primers used in Illumina amplification. A Visium Spatial Gene Expression library comprises standard Illumina paired-end constructs which begin and end with P5 and P7.Raw FASTQ files and histology images were processed (SpaceRanger software) for genome alignment. Raw output files for each group were read into R studio (Seurat 4.1.2). Normalization across spots was performed with the SCTransform function. The spatial cluster gene signature overlap correlation matrix was generated by first taking all genes differentially expressed (avg_log2FC > 1 and adjusted p value < 0.05) across all ST clusters. Pearson correlation was calculated across this matrix and hierarchical clustering of ST clusters was performed with the heatmap function with the ggplots package. Signature scoring derived from scRNA-seq and ST signatures was performed with the AddModuleScore function.

### Histopathology, Immunohistochemistry, and immunofluorescence microscopy

The samples were fixed with 4% paraformaldehyde at least 48 hours before ethanol and xylene dehydration. H&E staining and Masson’s trichrome staining were performed for the observation of re-epithelialization and collagen fiber deposition. Immunofluorescence staining for cytokeratin 5 (ab52635, Abcam, 1:200) and cytokeratin 10 (ab76318, Abcam, 1:150) and was performed for to assess keratinization. For the evaluation the infiltration of immune cells, immunohistochemistry staining for CD3 (14-0032-82, Thermo Fisher Scientific, 1:100), CD68 (ab283654, Abcam, 1:100), Ly6G (ab238132, Abcam, 1:2000) and immunofluorescent staining for CD3 (14-0032-82, Thermo Fisher Scientific, 1:100), CD68 (ab201844, Abcam, 1:250) were performed.

### Flow cytometry analysis

The surface markers of T cells (CD3), monocyte, and macrophages (CD11b, CD68) were examined using flow cytometry to quantification the infiltration of immune cells. The digested cell solutions were coincubated with antibodies against CD68 (Allophycocyanin, ab221251, Abcam), CD3 (fluorescein isothiocyanate, catalog no. 100203, BioLegend), and CD11b (PerCP/Cy5.5, catalog no. ab210299, Abcam) at 1:400 dilution for 30∼60 min at 4°C in the dark (100 µl per antibody per sample). Before the staining of CD3, cells were preincubated with purified anti-CD16/CD32 antibody (catalog no. 101301, BioLegend) (1.0 µg per 10^6^ cells in 100 µl volume) for 5 to 10 min to block Fc receptors. Flow cytometry analysis was performed using Attune Nxt flow cytometer (Thermo Fisher Scientific) and FlowJo 10.8.1. The experiments were performed three times independently (n = 3).

### Statistics and reproducibility

At least three independent replicates were conducted to obtain repeatable data, and the data was dealt with Case Viewer software, Image J software, and Prism 8.0 software. Statistical analyses were performed by Student’s t-test or one-way analysis of variance (ANOVA) with Tukey post-hoc test. The numerical data are presented as mean ± standard deviation. A value of p < 0.05 was considered statistically significant (*p < 0.05, **p < 0.01, ***p < 0.001), and ns means no statistically significant difference.

## Acknowledgments

We thank the Novogene company for RNA sequencing work, and the National Clinical Research Center for Oral Diseases & State Key Laboratory of Oral Diseases for animal experiment, and the Analytical & Testing Center of Sichuan University for the fabrication of biomaterials. We thank Chen Hu, Xin-hui Li, Jia-yu Gao and Sheng-an Rung for the help with analysis.

## Author contributions

Conceptualization: YY, CCY, YM, LL, YLQ

Methodology: CYC, LL, CBW

Investigation: YY

Visualization: YY

Writing – original draft: YY

Writing – review & editing: CCY

Project administration: CCY

Funding acquisition & Supervision: YM, YLQ

## Data and materials availability

All data to support the conclusions in this manuscript can be found in the figures and the supplementary materials. Any other data can be requested from the corresponding authors.

## Extended data

**Extended Data Figure 1.**
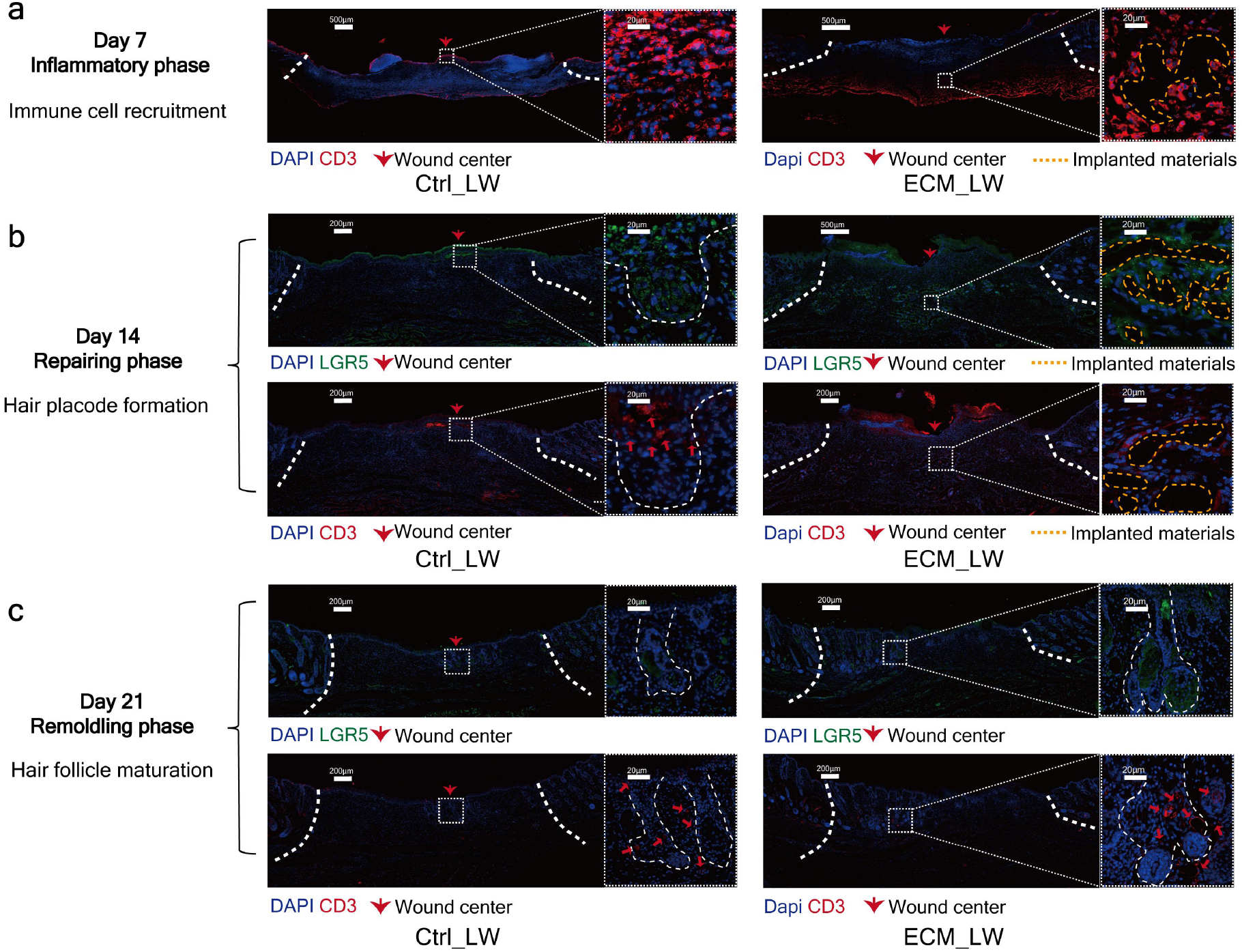
Evaluation the distribution of the HFSC and T cell. (a) Immunostaining for CD3 (red) showing the significant CD3^+^ T cell recruitment in biomaterial-implanted group on day 7. (b) Immunostaining for LGR5 (green) and CD3 (red) reveals LGR5^+^ HFSC and CD3^+^ T cell infiltration around biomaterials on day 14. (c) Immunostaining for LGR5 (green) and CD3 (red) reveals many LGR5^+^ HFSC within the bulge region of neogenic hair follicles adjacent to CD3^+^ T cells on day 21.

**Extended Data Figure 2.**
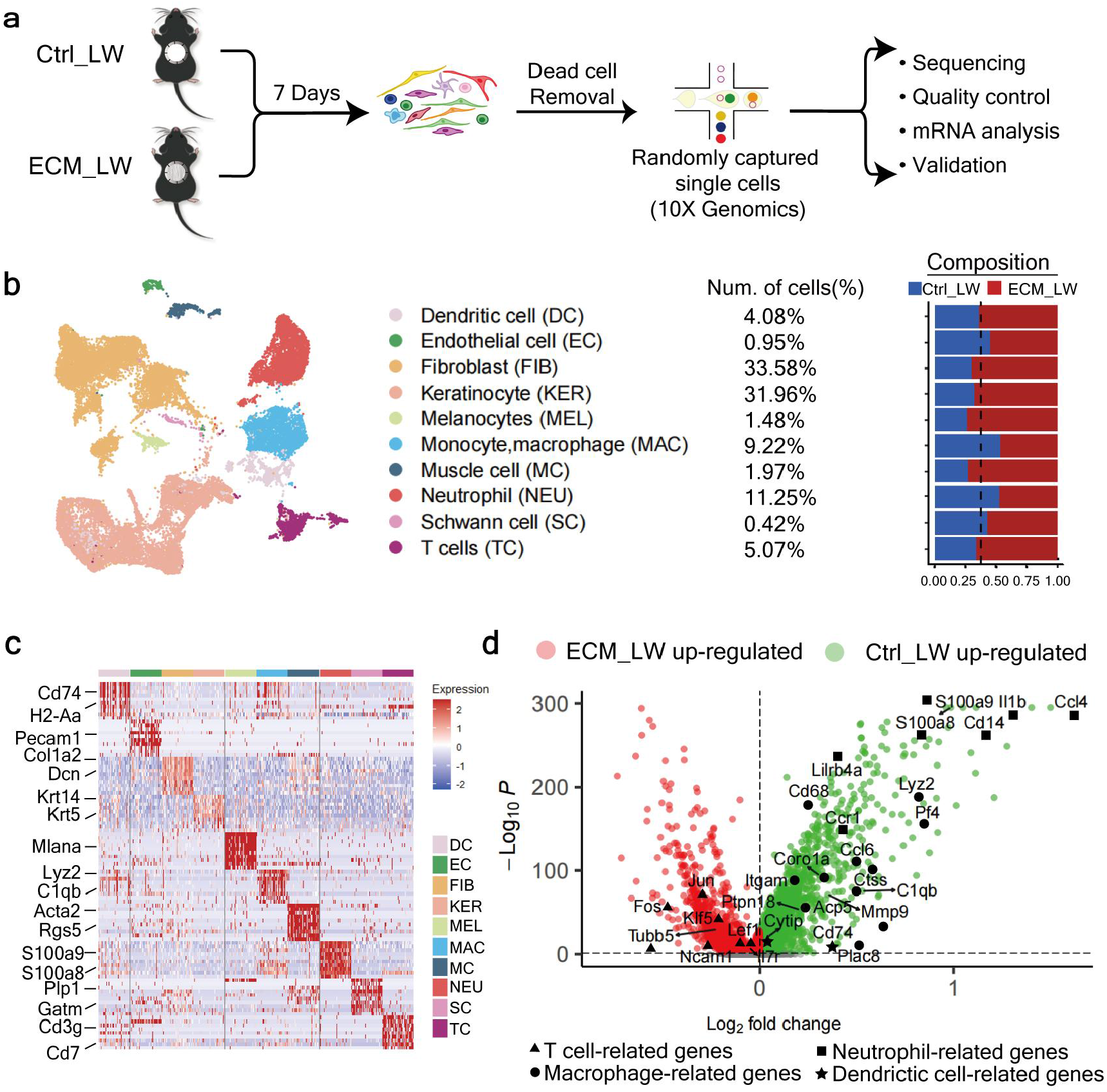
Overview of the single-cell transcriptome analysis between ECM_LW and Ctrl_LW. (a) Single-cell experiment workflow. (b) Cells are categorized into 10 main clusters. The number of cell populations in each cluster, number of cells (%), and composition of ECM_LW and Ctrl_LW groups are listed. (c) Heatmap showing top10 marker genes for each cluster. (d) Volcano plot showing interested differentially expressed genes in ECM_LW and Ctrl_LW group (scRNA-seq).

**Extended Data Figure 3.**
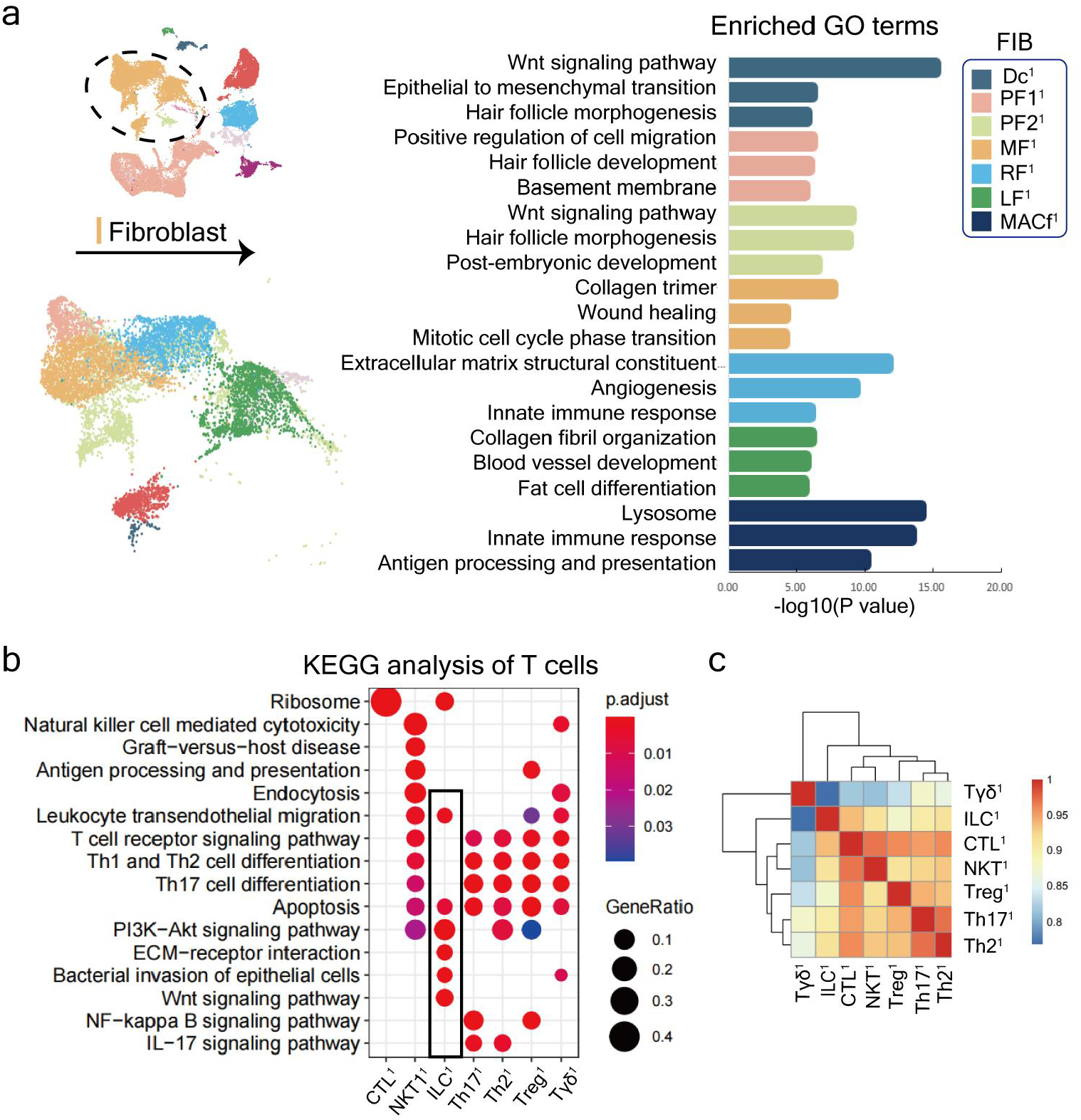
Further analysis of fibroblasts and T cells between ECM_LW and Ctrl_LW. (a) GO enrichment analysis of fibroblast subtypes. (b) KEGG enrichment analysis of T cell subtypes. (c) Pearson correlation analysis of T cell subsets based on average gene expression.

**Extended Data Figure 4.**
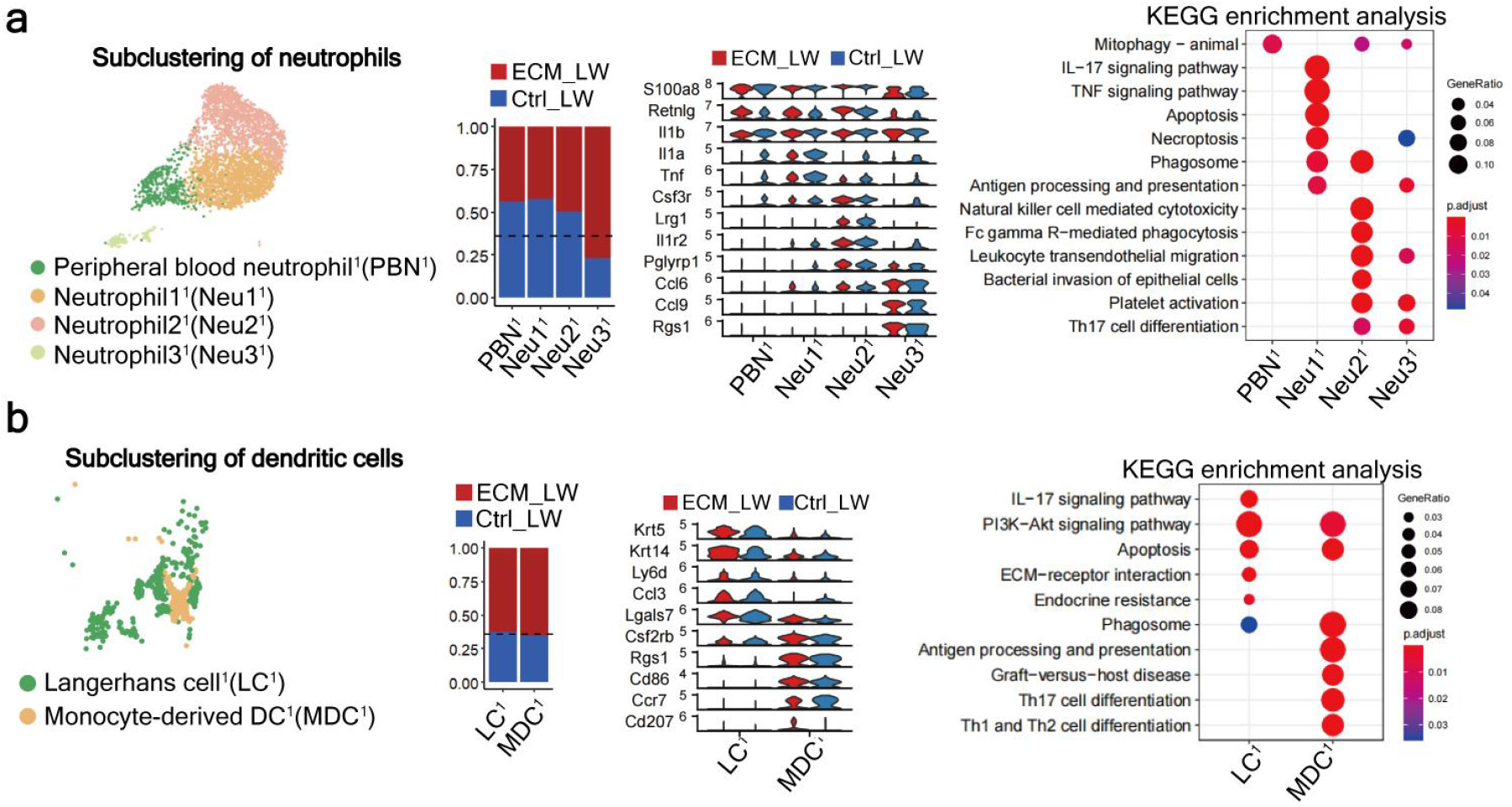
Subclustering analysis of neutrophils and dendritic cells between ECM_LW and Ctrl_LW. (a) Subclustering of neutrophils showing four subsets. The marker genes, composition and KEGG enrichment analysis for each subset are listed. (b) Subclustering of dendritic cells showing two subsets, marker genes, composition and KEGG enrichment analysis are listed.

**Extended Data Figure 5.**
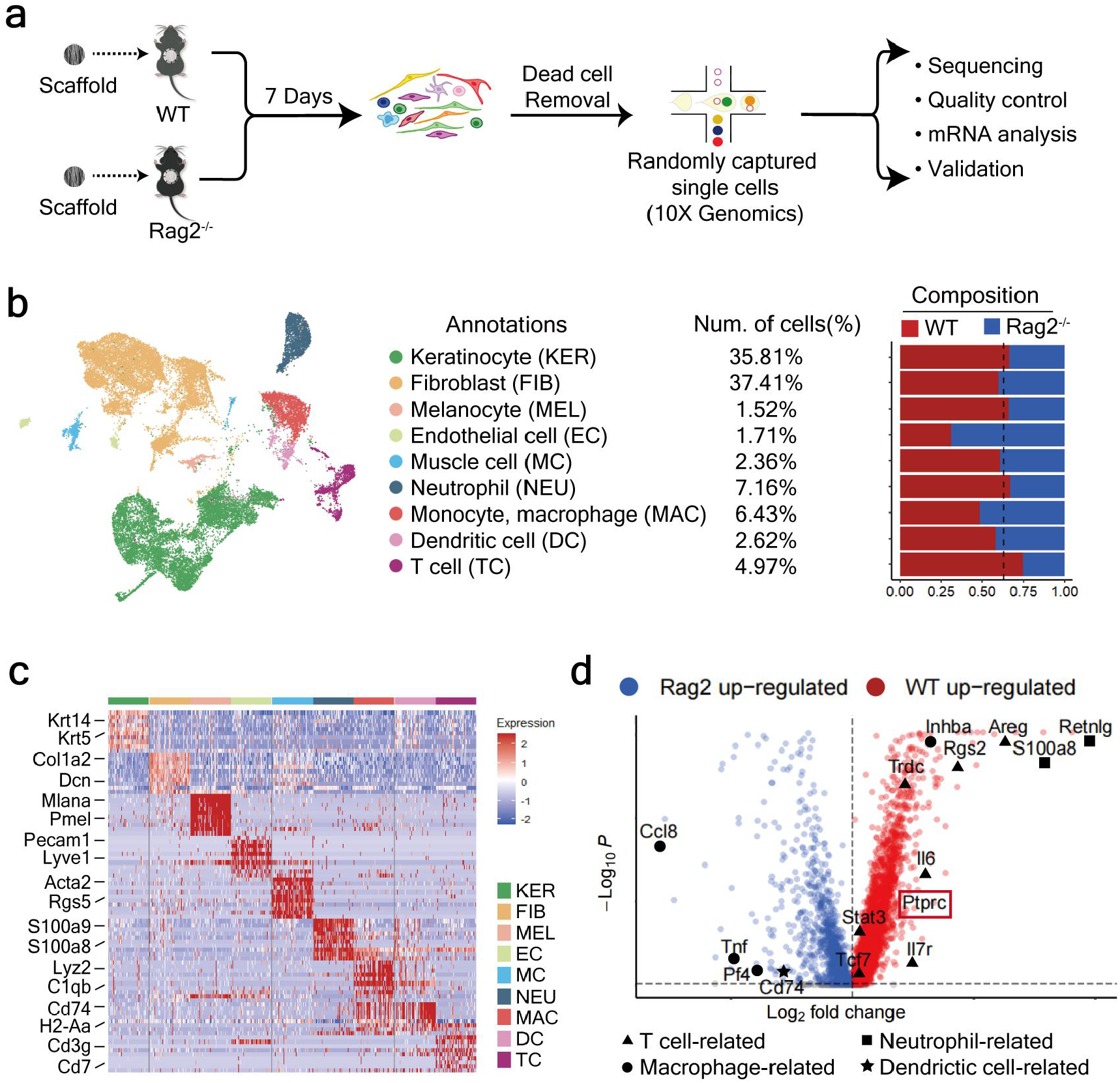
Overview of the single-cell transcriptome analysis between WT and Rag2^-/-^ mice treated with biomaterials. (a) Single-cell experiment workflow. (b) Cells are categorized into nine main clusters. The number of cell populations in each cluster, number of cells (%), and composition of ECM_LW and Ctrl_LW groups are listed. (C) Heatmap showing top10 marker genes for each cluster. (d) Volcano plot showing interested differentially expressed genes in WT and Rag2^-/-^ group (scRNA-seq).

**Extended Data Figure 6.**
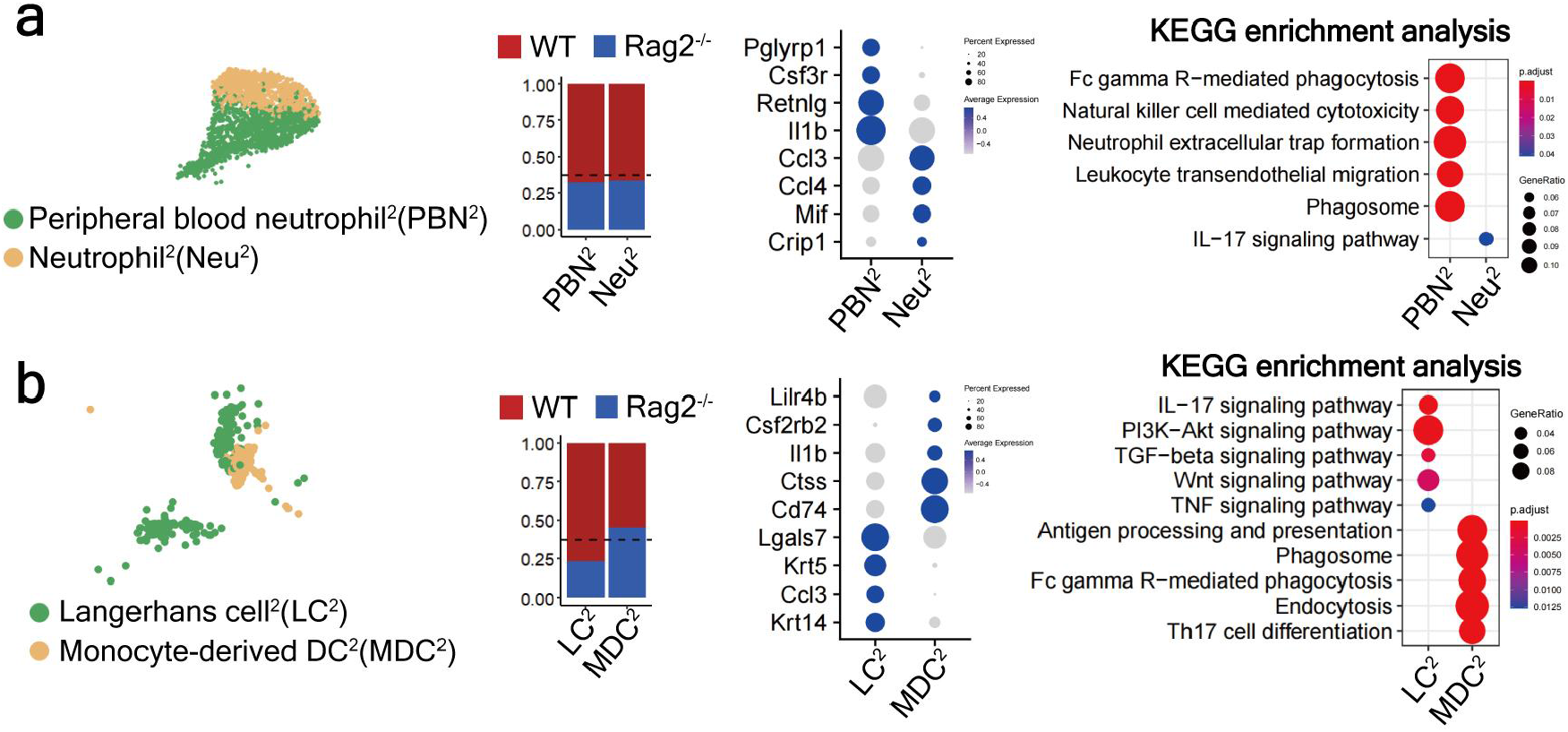
Subclustering analysis of neutrophils and dendritic cells between WT and Rag2^-/-^ mice. (a) Subclustering of neutrophils showing two subsets. The marker genes, composition and KEGG enrichment analysis for each subset are listed. (b) Subclustering of dendritic cells showing two subsets, marker genes, composition and KEGG enrichment analysis are listed.

**Extended Data Figure 7.**
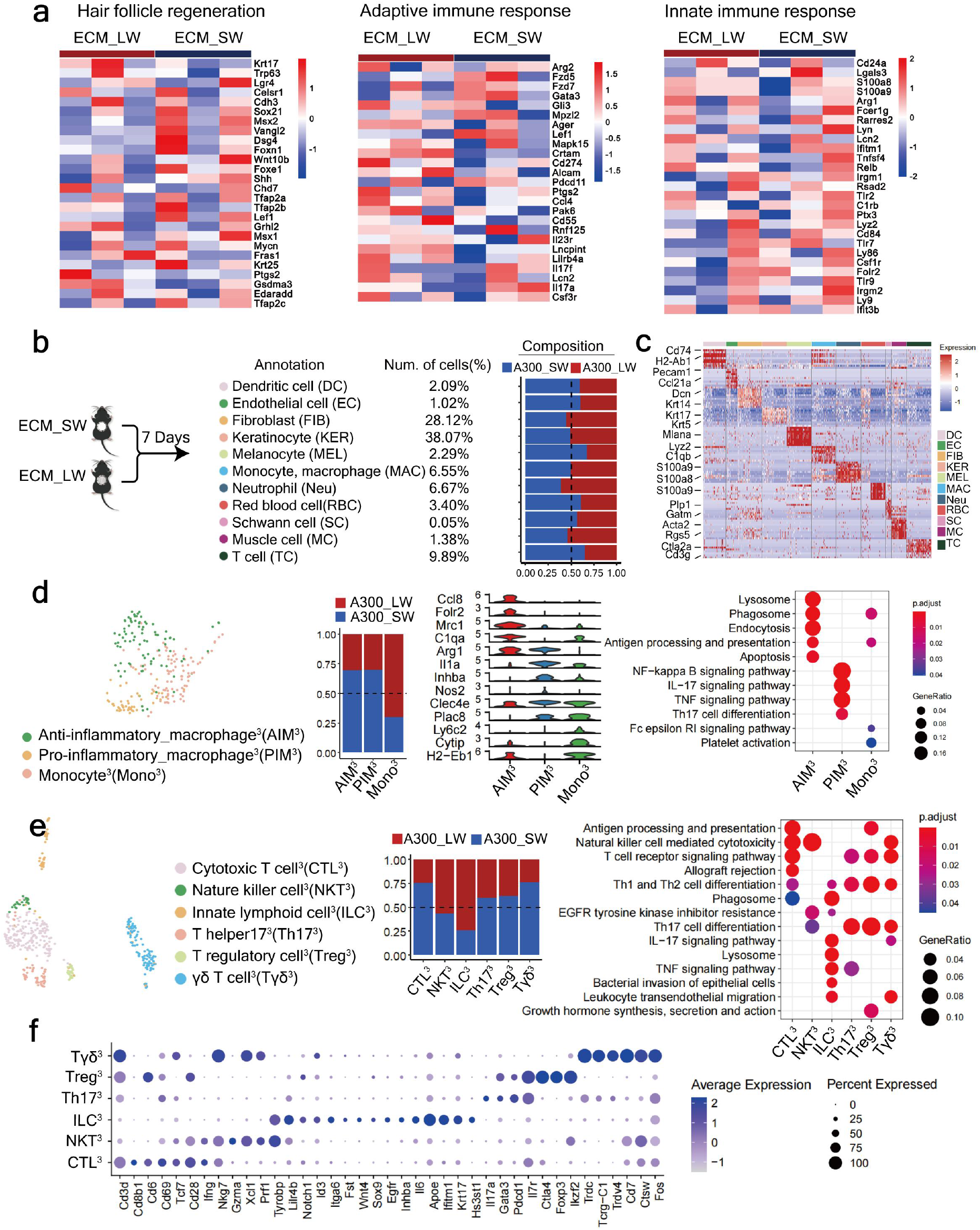
ScRNA-seq and ST analysis between ECM_SW and ECM_LW. (A) Heatmap showing expression of hair follicle regeneration, adaptive immune response, and innate immune response related genes (bulk seq). (B) Single cells are categorized into 11 main clusters. The number of cell populations in each cluster, number of cells (%), and composition of ECM_LW and ECM_SW groups are listed. (C) Heatmap showing top10 marker genes for each cluster. (D) Subclustering of MAC showing three subsets; The composition for each subset is listed; Marker genes for each MAC subset and enrichment analysis of KEGG pathway. (E) Subclustering of T cells showing seven subsets; The composition for each subset is listed and enrichment analysis of KEGG analysis. (F) Marker genes for T cell subsets.

